# When Metabolism Became Messaging: The Stepwise Evolution of the GABAergic Signaling System

**DOI:** 10.64898/2026.09.08.749565

**Authors:** Ankit Thakur, Mahesh Kulharia

## Abstract

The GABAergic system is the principal inhibitory signaling machinery in bilaterian nervous systems, yet its evolutionary assembly remains unresolved. Here, we reconstruct the origin and diversification of the complete GABAergic toolkit-including biosynthetic and catabolic enzymes, transporters, and ionotropic and metabotropic receptors-across 89 proteomes, representing 53 metazoan species and 36 outgroup lineages using phylogenomics and gene tree-species tree reconciliation. Our analyses reveal that GABAergic signaling did not emerge as a unified synaptic system but assembled stepwise from pre-existing metabolic components. Core enzymes of the GABA shunt (GAD1 and ABAT) predate nervous systems and are present in basal metazoans, indicating an ancestral metabolic or paracrine role. The emergence of GABA signaling in Cnidaria coincides with the recruitment of plasma membrane transporters and metabotropic GABA-B receptors, supporting early modulatory functions. In contrast, key components required for fast synaptic inhibition-vesicular transporter VIAAT and ionotropic GABA-A receptors-appear only in bilateria, marking a major functional transition. The asymmetric distribution of GABA-B subunits further suggests an ancestral promiscuous signaling state prior to obligate heterodimerization. Together, these findings support a three-stage evolutionary model in which inhibitory neurotransmission arose through progressive co-option and specialization of ancient molecular modules over ∼500-600 million years.

## 1. Introduction

Gamma-aminobutyric acid (GABA) is universally recognized as the principal inhibitory neurotransmitter in the central nervous system of bilaterians, where it is essential for regulating neuronal excitability and driving neural network homeostasis (Pierobon 2021). However, the evolutionary roots of the GABAergic signaling apparatus significantly predate the advent of centralized nervous systems, as evidenced by the widespread distribution of core biosynthetic enzymes, such as glutamate decarboxylase (GAD), across early-diverging, non-bilaterian metazoans. In the most basal, nerveless lineages such as Porifera (sponges), components of the GABAergic toolkit likely functioned as metabolic intermediates and paracrine signaling molecules to coordinate primitive, non-synaptic contractile behaviors (Elliott and Leys 2010). The subsequent functional transition of GABA from a diffuse metabolic modulator to a rapid, high-fidelity synaptic transmitter represents a stepwise evolutionary achievement, heavily dependent upon the lineage-specific expansion and structural rearrangement of specialized metabotropic and ionotropic receptors near the divergence of the cnidarian-bilaterian ancestor (Francis et al. 2017).

### 1.1 GABA Biosynthesis: The Dual Glutamate Decarboxylases

The genesis of GABAergic signaling primarily depends on the catalytic activity of glutamate decarboxylases (GADs), a class of pyridoxal-5’-phosphate (PLP)-dependent enzymes that catalyze the irreversible α-decarboxylation of L-glutamate to γ-aminobutyric acid and carbon dioxide (Martin and Rimvall 1993). In mammals, two distinct isoforms of this enzyme-GAD67 and GAD65, encoded respectively by the *GAD1* and *GAD2* genes located on human chromosomes 2 and 10-have emerged from an ancestral duplication event, each contributing uniquely to the cellular GABA economy. This functional divergence of GAD isoforms represents an important evolutionary specialization: GAD67, the product of *GAD1*, functions as the rate-limiting enzyme for bulk cytoplasmic GABA synthesis and maintains the ambient GABA pool required for metabolic and extrasynaptic signaling, whereas GAD65, encoded by *GAD2*, exhibits preferential association with synaptic vesicles and participates in the dynamic adjustment of neurotransmitter release during high-frequency neural activity (Soghomonian and Martin 1998). Both enzymes exist as homodimeric structures with three functionally distinct domains (PLP-binding, N-terminal, and C-terminal), and critically, dimerization is essential for catalytic activity, a structural constraint that has been preserved across evolution, with mutations disrupting optimal dimer association linked to disorders including schizophrenia.

### 1.2 GABA Catabolism: The GABA Shunt and Mitochondrial Integration

Following its synaptic activity, GABA is catabolized within mitochondria via the GABA shunt, a metabolic pathway that channels neurotransmitter-derived carbon skeleton into the tricarboxylic acid cycle and, simultaneously, constrains GABA accumulation to physiological ranges(Patel et al. 2005; Schousboe et al. 2013). This catabolic pathway is initiated by GABA transaminase (GABA-T), the product of the *ABAT* gene, a mitochondrial matrix enzyme that catalyzes the transamination of GABA to succinic semialdehyde (SSA) using α-ketoglutarate as an amino acceptor (Besse et al. 2015).

The subsequent step in GABA catabolism involves succinate-semialdehyde dehydrogenase (SSADH), encoded by *ALDH5A1*, which oxidizes SSA to succinate, a tricarboxylic acid (TCA) cycle intermediate, thereby linking GABA metabolism to cellular energy production(Chambliss et al. 1995).

### 1.3 GABA Reuptake: The Solute Carrier 6 Family Transporters

Termination of GABAergic neurotransmission depends upon the rapid removal of GABA from the synaptic cleft through the action of high-affinity membrane transporters belonging to the solute carrier 6 (SLC6) family (Kristensen et al. 2011). The mammalian GABAergic system employs four distinct GABA transporter isoforms-GAT1 (*SLC6A1*), GAT2 (*SLC6A13*), GAT3 (*SLC6A11*), and BGT1 (*SLC6A12*) whose tissue distributions and cellular localizations reflect a division of labor in regulating GABA homeostasis across different neural compartments. GAT1 (SLC6A1) functions as the principal GABA transporter throughout the central nervous system, expressed in both GABAergic neurons and astrocytes, where it mediates the sodium- and chloride-dependent reuptake of GABA down the electrochemical gradient, thereby establishing the fundamental temporal dynamics of GABAergic signaling (Bragina et al. 2008). This transporter comprises 12 transmembrane helices flanked by N- and C-terminal cytoplasmic domains, with three extracellular glycosylation sites in the second extracellular loop-architectural features conserved across diverse mammalian species and reflecting strong structural constraint (Bennett and Kanner 1997).

The structural heterogeneity of GABA transporters extends beyond their primary sequences to their physiological properties: GAT1 exhibits high affinity and selectivity for GABA and predominates in brain, spinal cord, and retina where rapid synaptic GABA reuptake is essential; GAT2 and BGT1, conversely, display broader substrate selectivity and lower abundance in the CNS, instead populating peripheral tissues including liver, kidney, and meninges where GABA has less defined synaptic roles.

A more recently characterized transporter, the vesicular inhibitory amino acid transporter (VIAAT, encoded by *SLC32A1*), localizes to synaptic vesicles and utilizes a proton electrochemical gradient to drive the active transport of newly synthesized GABA into vesicles for release-a compartmentalization that, combined with the plasma membrane reuptake activity of GATs, establishes a recycling cycle that ensures efficient reutilization of GABA and the sustained capacity for inhibitory neurotransmission (Zhou et al. 2012).

Despite their shared physiological role in regulating GABAergic tone, these transporters exhibit profound evolutionary and structural divergence that fundamentally dictates their distinct transport mechanisms. The plasma membrane transporters (GAT1, GAT2, GAT3, and BGT1) all belong to the SLC6 family and share a highly conserved LeuT-like structural scaffold characterized by 12 transmembrane (TM) helices (Motiwala et al. 2022). Within this SLC6 subgroup, structural diversity is primarily localized to specific amino acid substitutions within the central orthosteric binding pocket and conformational variations in the extracellular loops, which collectively govern their differential affinities for GABA versus broader substrates like betaine. In stark structural contrast, the vesicular transporter VIAAT belongs to the entirely distinct SLC32 family, completely lacking the LeuT-like architecture and instead possessing a unique topological fold comprising 9 to 10 transmembrane helices (Gasnier 2000). Consequently, while the structural core of the SLC6 GATs is optimized to harness extracellular sodium and chloride gradients for symport, the divergent structural architecture of VIAAT is specifically adapted to interface with the vesicular-type H^+^-ATPase, utilizing the vesicular proton electrochemical gradient for the active antiport loading of GABA.

### 1.4 GABA Signal Transduction: Ionotropic and Metabotropic Receptor Diversity

The final and functionally most diverse component of the GABAergic system comprises the GABA receptor superfamily-a complex array of proteins that translate chemical gradients into electrical inhibition. These receptors are broadly categorized into two distinct classes based on their transduction mechanism: the fast, ionotropic GABA-A receptors (a member of the Cys-loop ligand-gated ion channel superfamily) and the slow, metabotropic GABA-B receptors (a member of the Class C G-protein-coupled receptor family) (Tretter and Moss 2008).

The GABA-A receptor complex represents the primary site of rapid synaptic inhibition in the vertebrate brain. These receptors typically assemble as heteropentameric chloride channels. While the mammalian genome encodes nineteen discrete subunit genes, the majority of synaptic receptors follow a canonical stoichiometry of 2α:2β:1γ (Baumann et al. 2001). Our study specifically analyzes the evolutionary history of the core subunit families required for this functional assembly: the α (alpha) subfamily (GABRA1-6), which primarily determines ligand affinity and kinetics; the β (beta) subfamily (GABRB1-3), which, together with the α subunits, forms the orthosteric agonist binding site; and the γ (gamma) subfamily (GABRG1-3), which is essential for synaptic clustering and benzodiazepine sensitivity (Miller and Aricescu 2014).

Distinct from the classical α-β-γ isoforms are the bicuculline-insensitive receptors formed by the ρ (rho) subfamily (GABRR1–3). Historically termed GABA-C receptors, these subunits primarily form homomeric pentamers (or pseudo-heteromers) highly expressed in the retina. Unlike the widespread heteromeric GABA-A receptors, the ρ-type receptors exhibit distinct pharmacological profiles and non-desensitizing kinetics, representing a specialized evolutionary adaptation within the broader ionotropic lineage (Amin and Weiss 1994).

Complementing this ionotropic diversity are the metabotropic GABA-B receptors, which mediate prolonged inhibitory modulation. Unlike the pentameric GABA-A channels, GABA-B receptors function as obligate heterodimers composed of GABA-B1 (GABBR1) and GABA-B2 (GABBR2) subunits (Kaupmann et al. 1998). This dimerization is functionally critical: GABA- B1 contains the specific extracellular Venus flytrap domain responsible for orthosteric ligand binding, while GABA-B2 masks an endoplasmic reticulum retention signal to enable membrane trafficking and houses the intracellular loops necessary for G-protein coupling (Stewart et al. 2018). Upon activation, these receptors couple to G_i/o_ proteins to inhibit voltage-gated Ca^2+^ channels and activate inwardly rectifying K^+^ (GIRK) channels, thereby suppressing neurotransmitter release and inducing postsynaptic hyperpolarization(Bettler et al. 2004).

## 2. Materials and methods

We have investigated the evolution of GABAergic genes, employing computational methods to analyze proteins and enzymes crucial for GABA synthesis, transport, catabolism/regulation, and receptor function across diverse organisms. All proteomic and protein sequence data were obtained from publicly available databases such as NCBI, UniProt-KB, (see Supplementary File A and B). As this study is entirely computational and relies on pre-existing, publicly accessible data, it did not require ethics approval. The methodology followed in this study was taken from *Goulty et al, 2023, Nature communications* (Goulty et al. 2023).

### 2.1 Species Selection

Proteome data for the selected species were primarily collected from publicly available databases such as NCBI and UniProtKB, with additional data sourced from other repositories. To ensure a broad representation of species, we included organisms from diverse taxonomic groups, spanning the following Eukaryota (domain) supergroups: Amoebozoa, Excavata, Opisthokonta, Cnidaria, Placozoa, Porifera, Holozoa, Fungi, Archaeplastida, Glaucophyta, and Rhodophyta. This diverse selection allows for a comprehensive analysis of the GABAergic systems across eukaryotic lineages.

To assess the completeness of the proteome data, we utilized the BUSCO tool on 109 proteomes with the eukaryota_odb10 database containing 255 single-copy orthologs. BUSCO estimates the completeness and redundancy of genomic or proteomic data(Seppey et al. 2019). While BUSCO analysis provided valuable insights into the quality of the proteome data, we did not exclude all species with low BUSCO scores for single copy ortholog, instead we used complete scores and employed a threshold cutoff of ≥ 80. Eventually we ended up with 89 species of proteomes for further study.

### 2.2 Homology and Orthogroup Identification

We curated a comprehensive set of reference protein sequences; known to be involved in GABAe synthesis, degradation, transport, and signaling; collectively governing GABAe homeostasis across peripheral and central nervous tissues, to reconstruct the GABAergic systems across diverse eukaryotic lineages. Based on an extensive literature review and curated biological databases, we included in our study a total of 26 key protein sequences that represent the core molecular machinery of the GABAergic system. These were grouped into enzymes/proteins involved in the two groups for synthesis and degradation (each with one sequence), five groups for transmembrane GABA dependent signalling receptors - alpha (6), beta (3), gamma (3), rho (3), type-B receptor (2), and three groups for cellular transport and intracellular trafficking - three categories Sodium- and Chloride-dependent GABA Transporters (3 sequences), Sodium- and Chloride-dependent betaine Transporters (1 sequences) and Vesicular Inhibitory Amino Acid Transporters (1).

Annotated protein information was retrieved from the Kyoto Encyclopedia of Genes and Genomes (KEGG) database and supplemented with manually curated entries from the literature mining, and UniProtKB/Swiss-Prot database(Kanehisa et al. 2025). These curated reference sequences served as seed queries for homology searches across our dataset of 89 eukaryotic proteomes (69 Opisthokonta species). This approach allowed us to identify potential homologs and reconstruct orthogroups corresponding to GABAergic-related genes in both well- characterized and poorly annotated species; these served as blast queries sequences for later searches.

We employed PsiBlast based searches using the curated query sequences described above(Altschul et al. 1997). The homologues identified were identified by the PsiBlast (conducted with 3 iterations and an E-value threshold of 1e^-25^) and results retained for downstream analysis (see Supplementary File C).

The filtered sequences were then subjected to BLASTp analysis for functional annotation by querying against the Swiss-Prot database, and the top hit (if sequence coverage >80% and percent identity >40%)(Pearson 2013; Rost 1999; L. Li et al. 2003) from the Swiss-Prot results was used to assign putative functional identity(see Supplementary File D).

The BLASTP hits across all of the 89 species were sorted by species. The intra-species redundancy was removed using CD-HIT with a sequence identity threshold of 100% (-c 1.0) and default parameters (see Supplementary File E) (Huang et al. 2010). This step ensured that only unique protein sequences were retained for every species, thereby reducing redundancy in downstream analyses and minimizing redundancy induced computational bias during orthogroup inference.

We employed Broccoli, a phylogeny-aware orthology assignment tool that uses a combination of sequence similarity and gene tree-based inference to delineate orthologous groups (OGs) from the filtered protein dataset (see Supplementary File F) (Derelle et al. 2020). Broccoli was run using default parameters and the maximum likelihood method for gene tree construction, which improves orthology resolution in large, taxonomically diverse datasets.

The resulting OGs of interest i.e., those that potentially represented known components of the GABAergic system were further analyzed using InterProScan (see Supplementary File G) (Blum et al. 2025). This combination of automated annotation and expert manual curation ensured accurate functional classification of GABAergic-related protein families across a wide phylogenetic range.

We grouped all the identified sequences into four main datasets: one for receptors, one for transporters, one for GABA synthesis, and one for its degradation enzymes. For each of these four datasets, we used the orthogroups identified by Broccoli. Specifically, we selected orthogroups based on their InterProScan domain annotations: those annotated as receptor-like proteins were used for the receptor dataset, transporter-like for the transporter dataset, GABA synthesis-like for the synthesis dataset, and catabolic-like proteins for the catabolic enzyme dataset.

To validate structural features consistent with canonical GABAergic receptors, we analyzed all candidate sequences for transmembrane (TM) domains using Phobius (Käll et al. 2007). Because our dataset encompasses both ionotropic and metabotropic receptor classes, distinct structural criteria were applied for sequence retention. The ionotropic GABA-A receptors (comprising the α, β, γ, and ρ subunits) belong to the Cys-loop ligand-gated ion channel superfamily, where each subunit canonically possesses four transmembrane domains (4-TM)(Miller and Aricescu 2014). Conversely, the metabotropic GABA-B receptors (subunits B1 and B2) belong to the Class C G-protein-coupled receptor (GPCR) family, which is characterized by a canonical seven-transmembrane (7-TM) architecture (Pin et al. 2003; Mao et al. 2020). Therefore, to encompass the 4-TM topology indicative of ionotropic subunits, the 7-TM topology characteristic of metabotropic Class C GPCRs, and potential predictive variances, all candidate sequences exhibiting between 4 and 7 transmembrane domains were retained for downstream analysis. We then used CD-HIT (-c 0.8) to reduce redundancy while preserving sequence diversity within this receptor set.

### 2.3 Sequence clustering

To explore sequence space relationships across these four functional groups, we employed CLANS2 version:2.2.2 (CLuster ANalysis of Sequences), a tool that visualizes pairwise sequence similarity in 2D/3D using an all-vs-all BLAST-based clustering approach (Frickey and Lupas 2004). Each group of receptors, synthetic enzymes, catabolic enzymes, and transporters- was analyzed independently. CLANS2 was run with pairwise similarity thresholds (P-values) ranging from 1e^-15^ to 1e^-100^. At more permissive thresholds (e.g., 1e^-20^), clusters appeared larger and more interconnected, capturing distant homologies. As the stringency was increased (e.g., 1e^-100^), clusters resolved into more distinct groups, allowing finer discrimination of protein subfamilies and evolutionary lineages. Final values of thresholds were empirically determined based on cluster stability across stringency gradients.

Sequences that formed well-defined clusters in CLANS at specific stringency levels were retained for downstream functional and phylogenetic analysis (see Supplementary File H). For receptors, sequences that formed stable and coherent clusters at a P-value threshold of 1e^-27^ were selected (see figure 6D, see Supplementary File O_D_ for comparison).

Similarly, for biosynthetic enzymes, a P-value of 1e^-23^ was found to be optimal for separating functionally coherent clusters (see figure 6A, see Supplementary File O_A_ for comparison). For catabolic enzymes, a more stringent threshold of 1e^-58^ was used (see figure 6B, see Supplementary File O_B_ for comparison). For Transporter proteins threshold was 1e^-40^ (see figure 6C, see Supplementary File O_C_ for comparison). To assess the robustness of sequence exclusion decisions we performed repeated clustering by varying threshold values and examined sequences that appeared or disappeared across threshold levels for taxonomic identities.

### 2.4 Phylogenetic Tree Construction and Analysis

Based on the CLANS clustering results, only sequences that formed coherent and well-supported clusters were retained from each orthogroup (OG) for phylogenetic investigation. These OGs, corresponding to distinct GABAergic gene families (e.g., biosynthetic enzymes, receptors, catabolic enzymes, and transporters), were aligned independently using MAFFT (see Supplementary File I) (Katoh et al. 2019).

Post-alignment, positions with more than 70% gaps were removed using trimAl to eliminate poorly aligned and phylogenetically uninformative regions (Capella-Gutiérrez et al. 2009). This step was crucial for reducing noise and improving the accuracy of tree reconstruction.

Phylogenetic trees were then reconstructed using IQ-TREE2 (Minh et al. 2020), employing ModelFinder to automatically select the best-fit substitution model based on the Bayesian Information Criterion (BIC) and the resulting tree topologies were visualized, colored, and annotated using FigTree v1.4.5 (Jovanovic and Mikheyev 2019). To assess nodal support, 1,000 ultrafast bootstrap (UFBoot) (Hoang et al. 2018) replicates were calculated. Additionally, Transfer Bootstrap Expectation (TBE) (Zaharias et al. 2023) values were computed using 100 non-parametric bootstrap replicates to provide more robust support for deep or weakly supported nodes (see Supplementary File J).

### 2.5 Rogue taxa analysis

We employed two complementary methods: the t-index and the Leaf Stability Index (LSI) (Wilkinson 2006), to identify and eliminate unstable or rogue sequences that could distort phylogenetic inference.

The t-index, implemented in IQ-TREE2, evaluates how frequently a taxon changes its phylogenetic position across bootstrap replicates. High t-index values indicate topological instability, suggesting that the taxon may be a rogue element affecting overall tree resolution. We primarily relied on TBE-based t-index scores, and sequences with a t-index > 2 were classified as unstable.

The Leaf Stability Index (LSI) provides a complementary measure by estimating the frequency with which a taxon maintains a consistent phylogenetic placement across bootstrap trees. It is calculated using quartet frequencies, making it less sensitive to small topological rearrangements. LSI was computed using RogueNaRok (Aberer et al. 2013), with TBE-based trees. In this metric, lower LSI scores denote higher instability.

Taxa flagged as unstable by either method t-index > 2 or low LSI scores-were considered problematic. These rogue sequences were excluded to prevent topological artifacts and improve tree robustness. Pruning was carried out using the rnr-prune function of RogueNaRok (see Supplementary File K). This combined approach allowed us to refine our gene trees by minimizing the impact of highly unstable taxa, thereby enhancing both tree robustness and biological interpretability.

### 2.6 Gene-tree/species-tree reconciliation

To investigate the evolutionary history of GABAergic gene families in the context of species evolution, we performed gene-tree::species-tree reconciliation using GeneRax (Morel et al. 2020) (--si-strategy HYBRID -r UndatedDL), an advanced tool designed for inferring duplication, transfer, and loss (DTL) events.

We used the undated duplication-loss (UndatedDL) model (-r UndatedDL), which allows reconciliation of gene trees with a species tree without requiring branch length information in absolute time units. This approach is particularly useful for diverse datasets where molecular dating is either unavailable or unreliable. Maximum likelihood gene trees were generated by IQ- TREE2 for the unrooted gene tree input. Subsequently, rather than inferring a species tree *de novo*, we utilized the Open Tree of Life (OToL) (Hinchliff et al. 2015) to provide a robust phylogenetic framework for our analysis. The comprehensive species tree was pruned to include only the 89 taxa corresponding to the proteomes used in this study. By adopting this externally curated topology, we ensured that the species tree was independent of the gene trees generated during our orthology search, thereby eliminating potential circularity in the downstream reconciliation process (see Supplementary File L).

The substitution model for each orthogroup (OG) was specified based on the best-fit model determined during gene tree construction in IQ-TREE2 using the Bayesian Information Criterion (BIC) (Zhao et al. 2008).

The reconciliation output was visualized using ThirdKind (Penel et al. 2022) (see Supplementary File Ma/b), a tool designed to interpret and graphically represent gene duplication, loss, and transfer events in the context of species evolution. This facilitated clear identification of evolutionary dynamics underlying the distribution and diversification of GABAergic-related gene families.

### 2.7 Data Visualization

Schematic illustrations and data visualizations were prepared using standard graphic design tools. Figures 1-4 were created using the vector graphics editor Inkscape, and Figure 5 was created using CLANS2 and edited using Inkscape(“Scientific-Inkscape/CITATION.Cff at Main · Burghoff/Scientific-Inkscape,” n.d.).

**Fig. 1.**
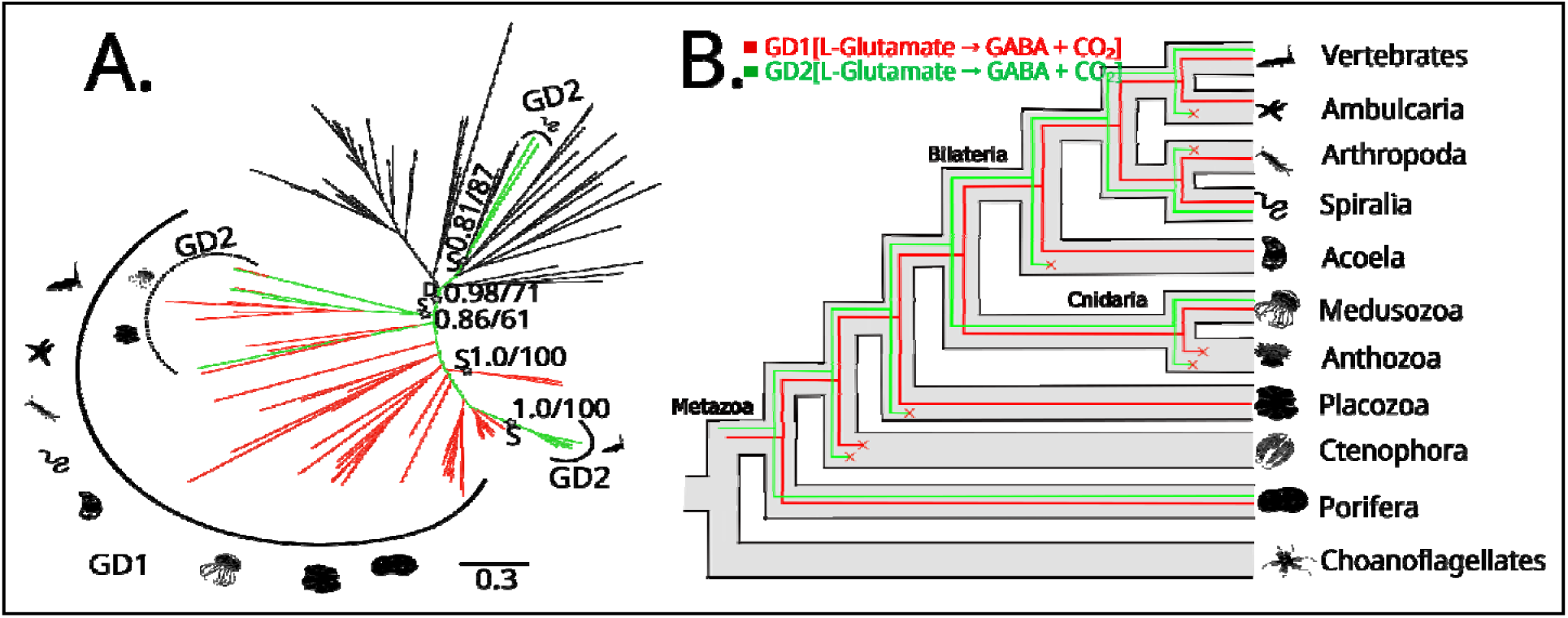
Phylogeny and reconciliation for Glutamate Decarboxylase 1 and 2 (GAD1 and GAD2). (A) Transfer bootstrap expectation tree and (B) simplified illustration of reconciliation calculated using Generax for GAD1 and GAD2 sequences. The nodal supports shown are transfer bootstrap expectation (TBE) scores (in decimal values), and ultrafast bootstrap proportion supports (in whole numbers) for key nodes. Asterisks (✰,if present) represents the Speciation (S) or Duplication (D) event with TBE and UFB support values. Dashed lines (if present) indicate sequences identified as unstable in t-index and leaf stability index (LSI) analysis. Silhouettes obtained from Phylopic.org.

**Fig. 2.**
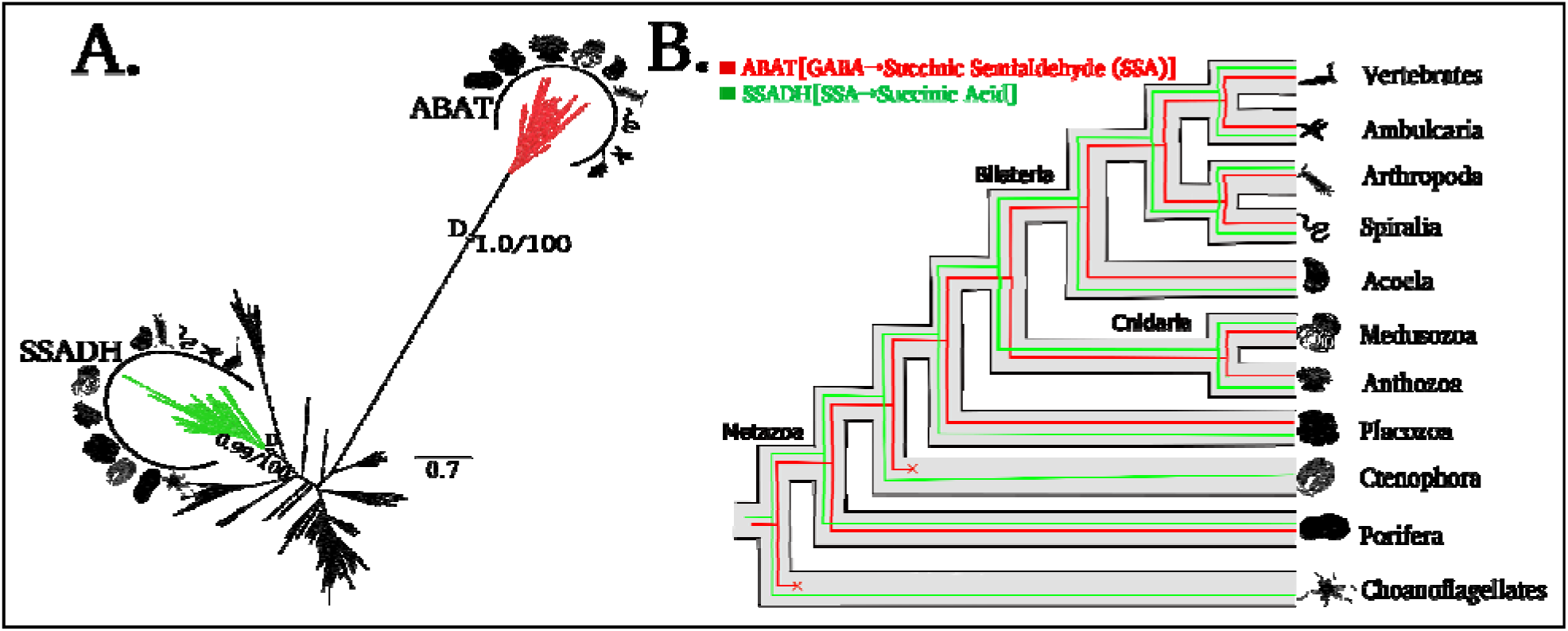
Phylogeny and reconciliation for 4-aminobutyrate aminotransferase (ABAT) and Succinate- semialdehyde dehydrogenase (SSADH). (A) Transfer bootstrap expectation tree and (B) simplified illustration of reconciliation calculated using Generax for ABAT and SSADH sequences. The nodal supports shown are transfer bootstrap expectation (TBE) scores (in decimal values), and ultrafast bootstrap proportion supports (in whole numbers) for key nodes. Asterisks (✰,if present) represents the Speciation (S) or Duplication (D) event with TBE and UFB support values. Dashed lines (if present) indicate sequences identified as unstable in t-index and leaf stability index (LSI) analysis. Silhouettes obtained from Phylopic.org.

## 3. Results

### 3.1 Evolutionary History of Glutamate Decarboxylase

We performed phylogenetic analysis of Glutamate Decarboxylase 1 and 2 (GAD1 and GAD2) to elucidate their evolutionary origins. Our results reveal that GAD1 forms a robust, expansive clade distributed widely across Metazoa, including basal lineages such as Porifera, Placozoa, and Medusozoa (Cnidaria), as well as Acoela, Spiralia, Arthropoda, Ambulacraria, and Vertebrata. Notably, GAD1 sequences were not detected in the Ctenophora dataset (see Figure 1A and 1B).

In contrast, GAD2 exhibited a more fragmented phylogenetic distribution, appearing in discrete clusters. One prominent GAD2 clade was restricted to Spiralia (TBE=0.81, UFB=87), while another highly supported subclade was exclusive to Vertebrata (TBE=1.0, UFB=100). Interestingly, several GAD2-like sequences from Medusozoa and Porifera clustered within the GAD1 clade. This phylogenetic positioning may represent ancestral or divergent forms related to GAD1-like sequences or early emergence events from a GAD1-like progenitor.

These findings indicate that neither GAD1 nor GAD2 is restricted to Bilateria; rather, both isoforms possess an ancient origin tracing back to the root of Metazoa. To further investigate the evolutionary drivers of these paralogs, we performed a gene-tree::species-tree reconciliation analysis. The results suggest that the divergence and subsequent expansion of these subclade were primarily driven by ancestral speciation events followed by lineage-specific duplications (see Supplementary File M_a/b_ and N_a/b_).

### 3.2 Evolutionary History of GABA Catabolic enzymes: ABAT and SSADH

We further investigated the evolutionary trajectory of the enzymes responsible for GABA degradation: 4-aminobutyrate aminotransferase (ABAT) and Succinate-semialdehyde dehydrogenase (SSADH). Phylogenetic reconstruction recovered robust monophyly for both ABAT (TBE=1.0, UFB=100) and SSADH (TBE=0.99, UFB=100), confirming their distinct evolutionary identities (see Figure 2A and 2B). Leaf Stability Index (LSI) and t-index analyses identified a single unstable sequence (of SSADH) from the demosponge *Sycon ciliatum*(Porifera), which was subsequently excluded to ensure the phylogenetic integrity of the final tree.

The taxonomic distribution of these enzymes revealed a striking asymmetry. ABAT sequence were identified across a broad range of metazoan lineages, including Porifera, Placozoa, Cnideria (Anthozoa and Medusozoa), Acoela, Spiralia, Arthropoda, Ambulacraria, and Vertebrata. However, similar to our observations for GAD1 and GAD2, ABAT was notably not detected in the Ctenophora dataset.

In contrast, SSADH (SSD) exhibited a near-ubiquitous distribution, being present in Choanoflagellates as well as all major metazoan classes, including Ctenophora.

Gene-tree::species-tree reconciliation analysis results suggest that the divergence and subsequent expansion of these clades were primarily driven by ancestral Duplication events (Supplementary File M_a/b_ and N_a/b_).

### 3.3 Evolutionary Dynamics of GABA transporters (SLC6 and SLC32 Families)

Efficient GABAergic signaling requires precise sequestration and transport, mediated by plasma membrane and vesicular transporters. We analyzed the evolution of Sodium- and Chloride- dependent GABA Transporters (GAT/SLC6 family) and Vesicular Inhibitory Amino Acid Transporters (VIAAT/SLC32 family).

Phylogenetic reconstruction recovered a highly supported monophyletic clade for VIAAT (TBE=1.0, UFB=100). Interestingly, VIAAT was found to be restricted to Bilateria, with sequences identified in Acoela, Spiralia, Arthropoda, Ambulacraria, and Vertebrata, but not detected in non-bilaterian lineages (see Figure 3A and 3B).

**Fig. 3.**
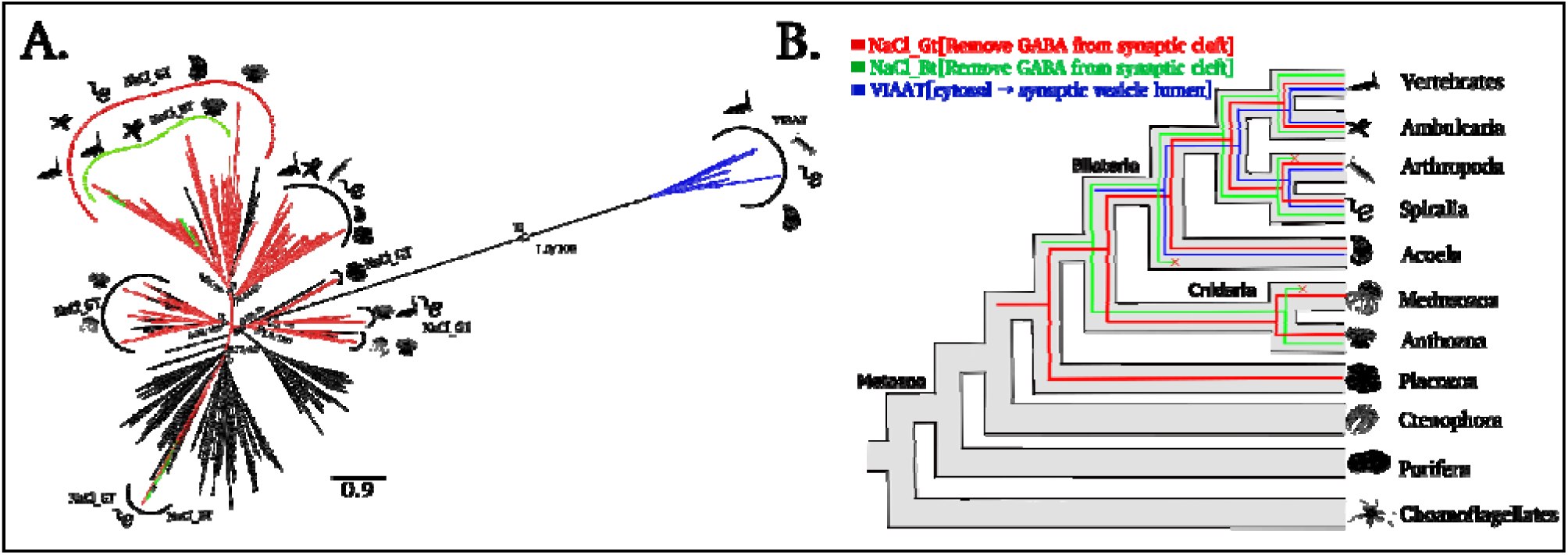
Phylogeny and reconciliation for Sodium- and Chloride-dependent GABA Transporters (GAT/SLC family) and Vesicular Inhibitory Amino Acid Transporters (VIAAT/SLC32 family). (A) Transfer bootstrap expectation tree and (B) simplified illustration of reconciliation calculated using Generax for GAT/SLC6 and SLC32 sequences. The nodal supports shown are transfer bootstrap expectation (TBE) scores (in decimal values), and ultrafast bootstrap proportion supports (in whole numbers) for key nodes. Asterisks (✰,if present) represents the Speciation (S) or Duplication (D) event with TBE and UFB support values. Dashed lines (if present) indicate sequences identified as unstable in t-index and leaf stability index (LSI) analysis. Silhouettes obtained from Phylopic.org.

**Fig. 4.**
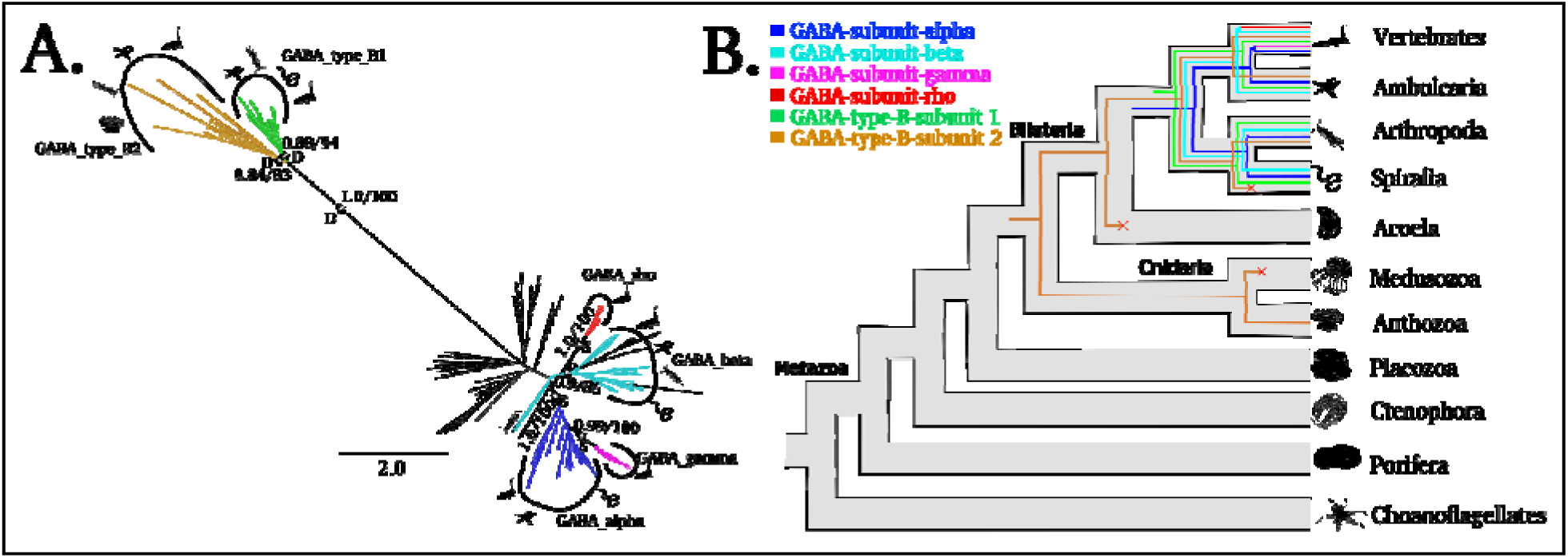
Phylogeny and reconciliation for the ionotropic GABA-A receptor subunits (α, β, γ, and ρ) and the metabotropic GABA-B receptor subunits. (A) Transfer bootstrap expectation tree and (B) simplified illustration of reconciliation calculated using Generax for GABA_A_, and GABA_B_ receptor sequences. The nodal supports shown are transfer bootstrap expectation (TBE) scores (in decimal values), and ultrafast bootstrap proportion supports (in whole numbers) for key nodes. Asterisks (✰,if present) represents the Speciation (S) or Duplication (D) event with TBE and UFB support values. Dashed lines (if present) indicate sequences identified as unstable in t-index and leaf stability index (LSI) analysis. Silhouettes obtained from Phylopic.org.

**Fig. 5.**
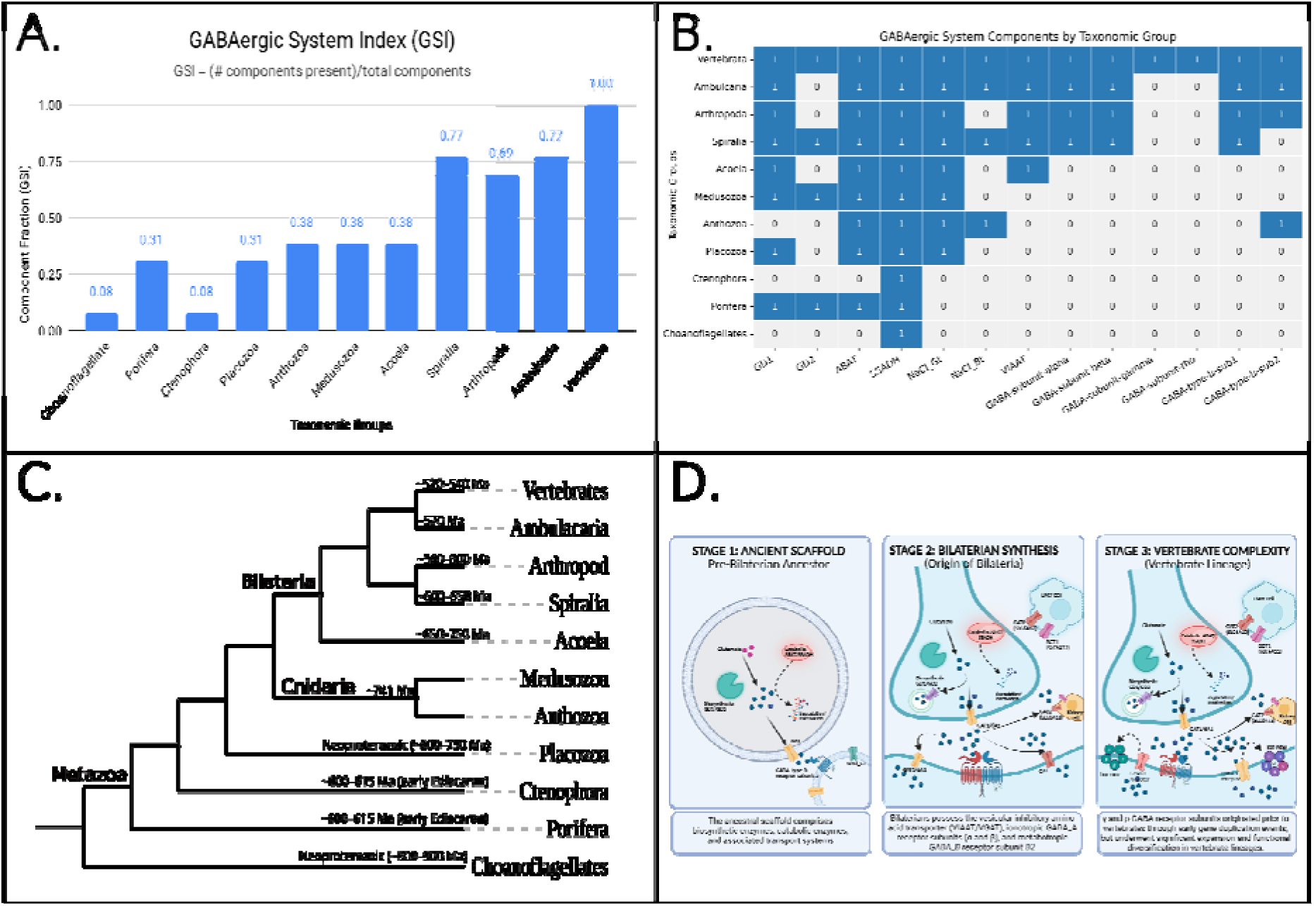
Evolutionary assembly and phylogenetic distribution of the GABAergic neurotransmitter system. **(A)** The GABAergic System Index (GSI) across major metazoan lineages. The bar chart calculates the fraction of total system components present in each taxonomic group, demonstrating the progressive evolutionary accumulation of the GABAergic toolkit from basal metazoans (e.g., Porifera, Placozoa) to its complete assembly in Vertebrata. **(B)** Presence-absence matrix detailing the phylogenetic distribution of 13 core GABAergic proteins across the analyze taxonomic groups. The matrix maps the origins of biosynthetic and catabolic enzymes (GAD1, GAD2, ABAT, SSADH), plasma membrane and vesicular transporters (NaCl_GT, NaCl_BT, VIAAT), and metabotropic/ionotropic receptor subunits (GABA-A α, β, γ, ρ; GABA-B R1, R2). Blue squares (1) denote the bioinformatic identification of an ortholog, while grey squares (0) indicate its absence. **(C)** Phylogenetic framework showing major metazoa lineages**. (D)** Schematic representation of the proposed Three-Stage Evolutionary Model for GABAergic signaling. **Stage 1 (Ancient Scaffold):** The pre-bilaterian emergence of the metabolic core, comprising ancient biosynthetic/catabolic enzymes and early transport systems. **Stage 2 (Bilaterian Synthesis/Innovation):** The establishment of rapid synaptic inhibition at the base of Bilateria, defined by the evolutionary innovation of the vesicular transporter VIAAT alongside ionotropic GABA-A (α and β) and metabotropic GABA-B (R1) receptor subunits. **Stage 3 (Vertebrate Complexity/Specialisation):** Lineage-specific structural and kinetic refinements driven by the vertebrate-specific expansion of the GABA-A γ and ρ subunits.

**Fig. 6.**
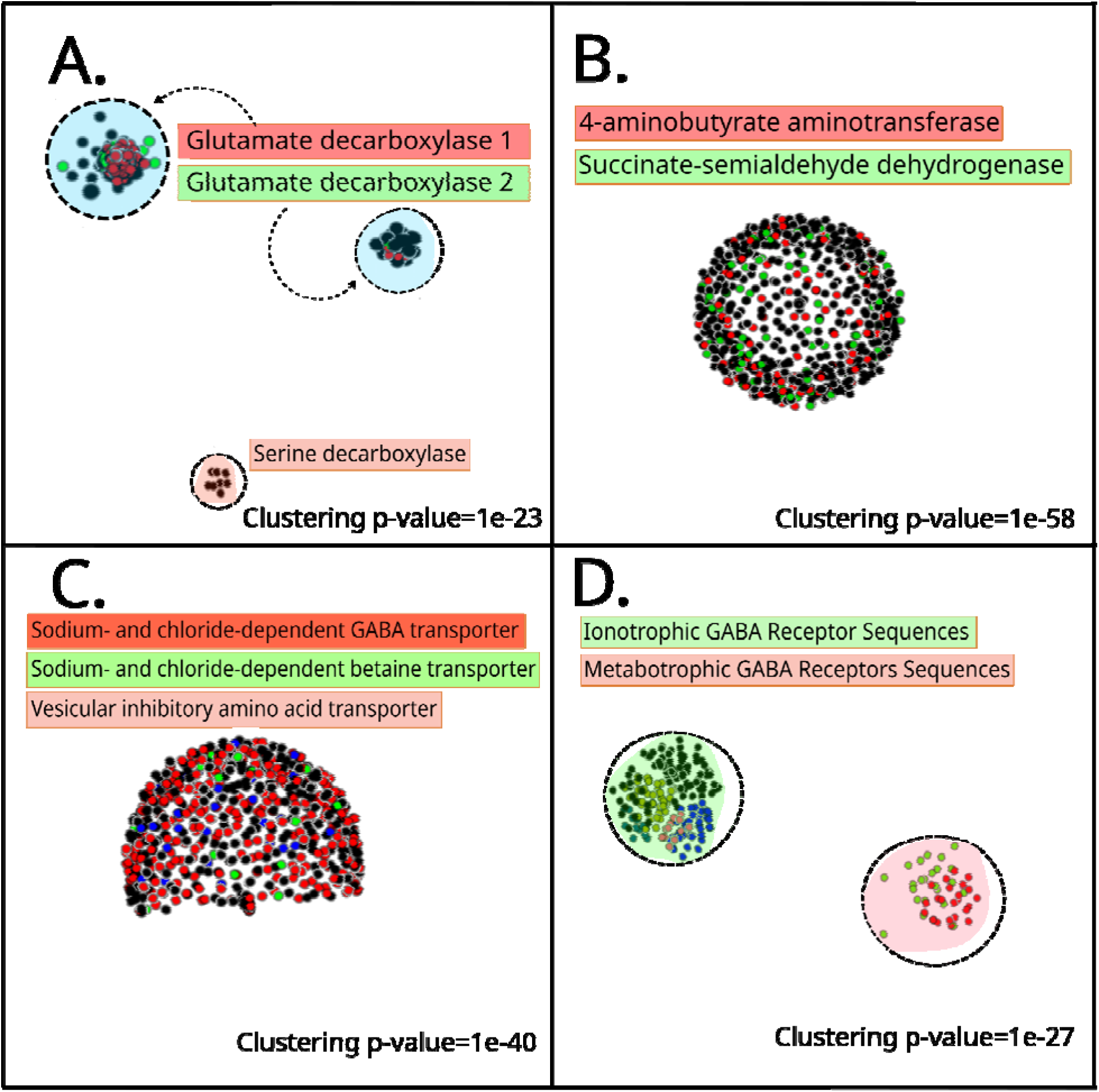
CLANS Analysis of the GABAergic Toolkit. CLANS (CLuster ANalysis of Sequences) plots illustrate the sequence similarity relationships across the four core functional groups of the GABAergic toolkit, with clustering driven by all-against-all BLAST P-values at specified stringencies (see Methodology). Each dot represents a single protein sequence, colored according to its functional annotation. **(A) Biosynthetic Enzymes** (P-value 1e-23). The plot displays two major, closely-associated clusters of Glutamate Decarboxylase (GAD) sequences. Both clusters contain sequences annotated as GAD1 (red dots) an GAD2 (green dots), indicating significant overlap and high sequence similarity between the two paralogous GAD isoforms. Sequences classified as other decarboxylases or those without definitive assignment are represented by black dots. **(B) Catabolic Enzymes** (P-value 1e-58). The analysis of the GABA catabolic enzymes reveals a single, continuous sphere-like structure, showing no clear distinction between the 4-aminobutyrate aminotransferase (ABAT, red dots) and Succinate-semialdehyde dehydrogenase (SSADH, green dots) sequences. This interminglin suggests a high degree of conserved domain architecture and sequence homology between these two sequential enzymes of the GABA shunt pathway. **(C) Transporters** (P-value 1e-40). The transporter sequences form a unified, hemisphere-like cluster. Sequences belonging to the plasma membrane Solute Carrier 6 (SLC6) family, specifically the GABA Transporters (NaCl_Gt, red) and Betaine Transporters (NaCl_Bt, green), are tightly grouped. Sequences from the structurally distinct Vesicular Inhibitory Amino Acid Transporter (VIAAT, lite brownish-red, SLC32 family) form a separate but physically associated region of the cluster, consistent with their shared role in GABA homeostasis despite profound structural divergence. **(D) Receptors** (P-value 1e-27). The plot shows two highly separated and independent clusters, reflecting the fundamental division of GABA receptors into two distinct structural superfamilies: **Metabotropic Cluster:** Contains the obligate heterodimer subunits of the Metabotropic GABA Type B receptor, GABABR1 (red) and GABABR2 (green). **Ionotropic Cluster:** Contains the components of the Cys-loop ligand-gated ion channel family, representing the fast Ionotropic GABA-A receptor subunits: alpha (α, blue), beta (β, olive), gamma (γ, pink), and rho (ρ, teal). The clear structural separation between these two clusters validates the evolutionary divergence of the Class C G-protein-coupled receptors (GABA-B) and the Cys- loop channels (GABA-A).

The plasma membrane transporters belonging to the SLC6 family exhibited a more complex evolutionary history. The GABA Transporter (GAT/NaCl_Gt) group was widely distributed across Placozoa, Cnidaria (Anthozoa and Medusozoa), and all major bilaterian clades. Within the GAT radiation, we identified specific subclades: one specialized to Cnidaria and another containing the Betaine/GABA Transporters (BGT/NaCl_Bt). BGT sequences were nested within larger GAT clades and were identified in Anthozoa, Spiralia, Ambulacraria, and Vertebrata (see Figure 3A).

To resolve the evolutionary history of these paralogs, we performed a gene-tree::species-tree reconciliation analysis. Our results indicate that the diversity of GAT (NaCl_Gt) sequences i primarily the result of extensive lineage-specific duplication events, particularly within Bilateria. The BGT (NaCl_Bt) isoform likely emerged through duplication within the GAT lineage but has likely undergone multiple independent loss events, leading to its absence in groups like Medusozoa and Arthropoda (see Supplementary File M_a/b_ and N_a/b_).

VIAAT similarly appears to be a product of a significant duplication event at the base of the Bilateria, potentially coinciding with the increased complexity of the bilaterian nervous system.

### 3.4 Evolutionary Diversification of GABA Receptors

To determine the emergence of GABA-sensing capabilities, we reconstructed the evolutionary history of both ionotropic and metabotropic GABA receptors. Our dataset included the ionotropic GABA-A receptor subunits (α, β, γ, and ρ) and the obligate heterodimeric metabotropic GABA-B receptor subunits (GABA-B1 and GABA-B2).

Phylogenetic analysis confirmed the presence of a core ionotropic signaling complex in ancestral bilaterians. The GABA-A α subunit (TBE=1.0, UFB=100) and β subunit (TBE=0.9, UFB=86) were identified across all major bilaterian clades, including Vertebrata, Ambulacraria, Arthropoda, and Spiralia (see Figure 4A). In contrast, the γ subunit (TBE=0.99, UFB=100) and ρ subunit (TBE=1.0, UFB=100) were found to be restricted exclusively to the Vertebrata lineage, suggesting these are later innovations or specific diversifications within the chordate radiation.

The metabotropic GABA-B receptor subunits exhibited a deeper, yet complex, evolutionary origin. Our phylogenetic reconstruction recovered GABA-B1 (TBE=0.88, UFB=94) and GABA- B2 (TBE=0.84, UFB=83) as distinct sister clades. During tree evaluation, t-index and Leaf Stability Index (LSI) analyses identified two unstable rogue sequences-one GABA-B1 subunit sequence from *Ciona intestinalis* and one GABA-B2 subunit sequence from *Branchiostoma belcheri*-which were subsequently excluded to ensure topological robustness. The taxonomic distribution of the retained sequences was found to be asymmetric: GABBR2 was identified in Anthozoa (Cnidaria) but GABBR1 was not detected, while GABBR1 was present in Spiralia but GABBR2 was notably not detected in the analyzed proteomes. The GABBR1 subunit was otherwise present across Vertebrata, Ambulacraria, Arthropoda, and the GABBR2 subunit in Vertebrata, Ambulacraria, and a single species of Arthropoda (*Daphnia pulex*). Both subunits were not detected in our Acoela dataset. This asymmetric, non-obligate distribution suggests that ancestral components of metabotropic GABA signaling predate the divergence of Bilateria and Cnidaria, arising from an ancestral gene duplication event prior to this split.

We employed gene-tree::species-tree reconciliation to resolve the mechanisms driving the expansion of this receptor toolkit:

Speciation-Driven Divergence: The diversification of the GABA-A α, γ, and ρ subunits appears to be primarily driven by speciation events, followed by lineage-specific retention in vertebrates. This suggests these subunits may have specialized roles that became fixed during the early evolution of the vertebrate nervous system.

Duplication-Driven Expansion: Conversely, both the GABA-A β subunit and the metabotropic GABA-B lineages show clear signatures of ancestral gene duplication events. Notably, reconciliation analysis revealed significant duplication events at the basal nodes of both the GABA-B1 and GABA-B2 sister clades(see Supplementary File M_a/b_ and N_a/b_). The initial duplication giving rise to these distinct B1 and B2 lineages likely occurred prior to the Cnidaria-Bilateria split, facilitating the evolution of the obligate heterodimer. Meanwhile, the β subunit expansion highlights a key duplication event that bolstered ionotropic receptor diversity in ancestral bilaterians.

## 4. Discussion

### The Stepwise Assembly of the GABAergic System

Our phylogenomic reconstruction reveals that the GABAergic system did not emerge as a monolithic functional unit but rather was assembled in distinct evolutionary stages with the progressive evolutionary accumulation of the GABAergic toolkit from basal metazoans (Fig. 5A and 5B). This "stepwise assembly" model challenges earlier assumptions that neurotransmitter systems evolved simultaneously with the first neurons(Najle et al. 2023). Instead, we observe a clear progression in complexity from metabolic capability to complex synaptic signaling.

### 4.1. The Ancient Metabolic Core (Porifera and Placozoa)

We have identified the core enzymes for GABA synthesis (GAD1) and catabolism (ABAT) in non-neural lineages such as Porifera (sponges) and Placozoa. This presence in organisms lacking a nervous system suggests that GABA may have originally functioned as a metabolic intermediate or a paracrine signaling molecule rather than a synaptic transmitter (Jékely 2021). Previous studies utilizing immunohistochemistry have reported GABA-like immunoreactivity in sponges, specifically in the marine demosponge *Chondrilla nucula* (Ramoino et al. 2007). Our genomic analysis supports the presence of the genetic machinery (GAD1, ABAT) definitively, validating that these organisms possess the intrinsic capacity to synthesize and degrade GABA, while other wet lab studies have demonstrated that GABA can induce coordinated contractions in sponges like *Tethya wilhelma*(Ellwanger et al. 2007; Ellwanger and Nickel 2006).

### 4.2. The Ctenophore Anomaly: Evidence for Independent Evolution?

A striking finding of our study is the apparent absence or strong divergence of GAD, ABAT, and GABA-specific transporters in the analyzed Ctenophora dataset, despite the presence of the universal enzyme SSADH. This corroborates the earlier genomic work by Moroz et al. (2014), which proposed that ctenophores evolved a "convergent" nervous system largely devoid of classical neurotransmitters (Moroz et al. 2014). This absence is highly significant for two reasons: **a)** it directly contradicts the expected toolkit if GABA were a classical, regulated neurotransmitter in Ctenophores, and **b)** it highlights a major evolutionary divergence. If GABA were a critical signaling molecule, the full complement of machinery for its synthesis (GAD), vesicular packaging (VIAAT), and reuptake (GAT) would be expected. The presence of only SSADH, a ubiquitous mitochondrial enzyme that is part of the GABA shunt but is also involved in general cellular metabolism, suggests that any low-level GABA in Ctenophora is likely a metabolic intermediate rather than a regulated synaptic signal. This finding, combined with the absence of other classical neurotransmitter machinery in their genome, is consistent with hypotheses proposing independent or divergent evolution of the ctenophore nervous system, relying on a fundamentally different chemical language than the Eumetazoa (Cnidaria + Bilateria)(Moroz et al. 2014).

### 4.3. The Emergence of Signaling: Metabotropic vs. Ionotropic (Cnidaria vs. Bilateria)

Our receptor analysis highlights a crucial evolutionary pivot. We detected the GABA-B2 subunit of the metabotropic GABA-B receptor in Anthozoa (Cnidaria), suggesting that "slow" G-protein coupled signaling predates the "fast" ionotropic transmission mediated by GABA-A receptors. This finding aligns with the "chemical brain" hypothesis, where slow, diffusive modulation evolved before precise, millisecond-scale synaptic transmission (Jékely 2021).

Previous pharmacological profiling of *Hydra* (Cnidaria) indicated sensitivity to GABA(Pierobon 2021; Pierobon et al. 1995). Our data clarifies this mechanism: Cnidarians likely utilize a system comprising GAT transporters and the GABA-B2 subunit for modulatory signaling. However, as they lack the GABA-B1 subunit (which is required for the obligate heterodimer), the specialized vesicular machinery (VIAAT) and ionotropic receptors found in Bilateria, their GABA dependent signaling is sub-optimal.

Our phylogenetic reconstruction reveals a striking asymmetrical distribution of metabotropic GABA-B subunits across early-diverging lineagesAsterisks such as the isolated presence of GABBR2 in Anthozoa and GABBR1 in Spiralia and Ctenophora. While this evolutionary decoupling challenges the canonical obligate heterodimer paradigm, it strongly aligns with emerging pharmacological and developmental evidence of independent subunit functionality. For instance, GABA-B2 can traffic to the cell surface independently, while GABA-B1 can circumvent the heterodimer constraint by forming functional complexes with other Class C GPCRs, such as the Calcium-Sensing Receptor (CaSR)(Chang et al. 2020). Furthermore, the robust expression of GABA-B1 prior to GABA-B2 during early mammalian embryonic development suggests a distinct, independent neurotrophic role that predates mature synaptic transmission (S. p. Li et al. 2006).

Therefore, our evolutionary data proposes a "Promiscuous Ancestor" hypothesis: early in metazoan evolution, GABA-B1 and GABA-B2 likely functioned either as homodimers or partnered promiscuously with other ancestral Class C GPCRs. The ancestral gene duplication event, as suggested by our reconciliation, provided the foundation for the eventual heterodimerization mechanism. The strict, obligate B1/B2 heterodimer observed in modern vertebrates is likely a later evolutionary refinement for highly specialized synaptic inhibition, rather than an ancestral prerequisite for metabotropic GABA sensing.

### 4.4. The Bilaterian Innovation: High-Speed Synaptic Transmission

The restriction of VIAAT and GABA-A subunits (α, β) to Bilateria marks the inferred emergence of the molecular machinery required for the modern inhibitory synapse. The co-emergence of a dedicated vesicular loader (VIAAT) and a fast ligand-gated ion channel (GABA- A) would have allowed for the temporal precision required for complex locomotion and behavior. Furthermore, our observation that GABA-A γ and ρ subunits are Vertebrata-specific suggests a late-stage refinement of this system. The addition of the γ subunit, known to anchor receptors at the synapse and modulate kinetics, likely facilitated the sophisticated neural processing seen in vertebrates.

### Limitations

A limitation of this study is the reliance on currently available proteomes, which may be incomplete or biased toward well-annotated species. Absence of genes should therefore be interpreted cautiously.

## 5. Conclusion

In this study, we provide a unified evolutionary history of the GABAergic system, reconciling the origins of synthesis, transport, and reception.

We propose a three-stage evolutionary model (see Fig. 5D):

**Stage A:** The Metabolic Stage (Pre-Neural/Porifera): GABA likely functioned initially as a metabolic intermediate via the GABA shunt (GAD1 + ABAT), serving non-synaptic roles in primitive metazoans(Elliott and Leys 2010).

**Stage B:** The Modulatory Stage (Cnidaria/Eumetazoa): The recruitment of plasma membrane transporters (GAT) and the GABA-B2 subunit allowed GABA to function as a slow, diffusive modulator in the early nervous systems of Cnidaria, in the absence of the VIAAT vesicular loading system(Pierobon 2021). Furthermore, the asymmetrical presence of single GABA-B subunits in basal lineages reinforces the "Promiscuous Ancestor" hypothesis, suggesting early metabotropic receptors functioned independently or with alternative GPCR partners prior to the evolution of the strict obligate heterodimer(Francis et al. 2017).

**Stage C:** The Specialized Synaptic Stage (Bilateria): The innovation of vesicular packaging (VIAAT) and fast ionotropic receptors (GABA-A α, β) enabled precise, rapid synaptic inhibition. This system underwent further refinement in Vertebrata through the duplication-driven emergence of specialized receptor subunits (γ, ρ). Genomic evidence strongly supports this vertebrate-specific expansion; as demonstrated by Tsang et al. (2006), All vertebrate [GABA-A receptor-like] gene sets in the study but no invertebrate ones exhibit the extensive and conserved pattern of gene clustering, indicating that the gene clusters were established early in vertebrate evolution. The authors further note that the modern receptor diversity could have arisen from an ancestral β-α-γ subunit cluster through 2 rounds of whole-genome duplication (WGD) early in vertebrate evolution (Tsang et al. 2007).

## The Ctenophore Anomaly and Independent Neural Evolution

The apparent absence or divergence of core GABAergic machinery in Ctenophora is consistent with the hypothesis of independent neural evolution in this clade(“The Ctenophore Genome and the Evolutionary Origins of Neural Systems | Nature,” n.d.). While ctenophores possess highly complex neural nets, muscles, and coordinated behaviors, extensive phylogenomic and metabolomic analyses have consistently failed to detect classical bilaterian neurotransmitters (including GABA, serotonin, dopamine, and acetylcholine) or their canonical receptors(Moroz 2015). Instead, ctenophores appear to utilize a completely distinct, independently derived chemical language-likely dominated by an extensive repertoire of unique, clade-specific neuropeptides(Moroz and Kohn 2016). This "alien" neurobiology implies that the molecular functionalization of the nervous system occurred at least twice during metazoan evolution, with the ctenophore lineage entirely bypassing the GABAergic synaptic toolkit used by Cnidaria and Bilateria.

## Acknowledgements

The authors acknowledge the University Grants Commission (UGC) and the Central University of Himachal Pradesh (CUHP) for providing institutional support, computational facilities, and Non-NET fellowship support.

## CRediT authorship contribution statement

MK conceptualized the work and AT carried out the research. The results were analyzed and manuscript written by AT and MK.

## Ethics approval and consent to participate

Not applicable. This study was entirely computational and used only publicly available proteomic and protein sequence data; no human participants, animals, or identifiable private data were involved, so ethics approval and consent to participate were not required.

## Data availability

The supplementary information associated with this manuscript is publicly available. The supplementary materials, including detailed datasets, additional information, and analyses, can be accessed via the following link: https://doi.org/10.6084/m9.figshare.31857025

These materials are provided to ensure transparency, and to facilitate further exploration of the results presented in this study.

## References

1. Aberer, Andre J., Denis Krompass, and Alexandros Stamatakis. 2013. “Pruning Rogue Taxa Improves Phylogenetic Accuracy: An Efficient Algorithm and Webservice.” Systematic Biology 62 (1): 162–66. 10.1093/sysbio/sys078.

2. Altschul, Stephen F., Thomas L. Madden, Alejandro A. Schäffer, et al. 1997. “Gapped BLAST and PSI-BLAST: A New Generation of Protein Database Search Programs.” Nucleic Acids Research 25 (17): 3389–402. 10.1093/nar/25.17.3389.

3. Amin, Jahanshah, and D. Weiss. 1994. “Homomeric Rho 1 GABA Channels: Activation Properties and Domains.” Receptors & Channels. https://www.semanticscholar.org/paper/Homomeric-rho-1-GABA-channels%3A-activation-and-Amin-Weiss/a237a945be8c32d999d0998a1063d509bdf3e667.

4. Baumann, Sabine W., Roland Baur, and Erwin Sigel. 2001. “Subunit Arrangement of γ- Aminobutyric Acid Type A Receptors*.” Journal of Biological Chemistry 276 (39): 36275– 80. 10.1074/jbc.M105240200.

5. Bennett, Estelle R., and Baruch I. Kanner. 1997. “The Membrane Topology of GAT-1, a (Na++ Cl−)-Coupled γ-Aminobutyric Acid Transporter from Rat Brain*.” Journal of Biological Chemistry 272 (2): 1203–10. 10.1074/jbc.272.2.1203.

6. Besse, Arnaud, Ping Wu, Francesco Bruni, et al. 2015. “The GABA Transaminase, ABAT, Is Essential for Mitochondrial Nucleoside Metabolism.” Cell Metabolism 21 (3): 417–27. 10.1016/j.cmet.2015.02.008.

7. Bettler, Bernhard, Klemens Kaupmann, Johannes Mosbacher, and Martin Gassmann. 2004. “Molecular Structure and Physiological Functions of GABA_B_ Receptors.” Physiological Reviews 84 (3): 835–67. 10.1152/physrev.00036.2003.

8. Blum, Matthias, Antonina Andreeva, Laise Cavalcanti Florentino, et al. 2025. “InterPro: The Protein Sequence Classification Resource in 2025.” Nucleic Acids Research 53 (D1): D444–56. 10.1093/nar/gkae1082.

9. Bragina, Luca, Ivan Marchionni, Azar Omrani, et al. 2008. “GAT-1 Regulates Both Tonic and Phasic GABAA Receptor-Mediated Inhibition in the Cerebral Cortex.” Journal of Neurochemistry 105 (5): 1781–93. 10.1111/j.1471-4159.2008.05273.x.

10. Capella-Gutiérrez, Salvador, José M. Silla-Martínez, and Toni Gabaldón. 2009. “trimAl: A Tool for Automated Alignment Trimming in Large-Scale Phylogenetic Analyses.” Bioinformatics 25 (15): 1972–73. 10.1093/bioinformatics/btp348.

11. Chambliss, Ken L., Deborah L. Caudle, Debra D. Hinson, et al. 1995. “Molecular Cloning of the Mature NAD+-Dependent Succinic Semialdehyde Dehydrogenase from Rat and Human.” Journal of Biological Chemistry 270 (1): 461–67. 10.1074/jbc.270.1.461.

12. Chang, Wenhan, Chia-Ling Tu, Frederic G. Jean-Alphonse, et al. 2020. “PTH Hypersecretion Triggered by a GABAB1 and Ca2+-Sensing Receptor Heterocomplex in Hyperparathyroidism.” Nature Metabolism 2 (3): 243–55. 10.1038/s42255-020-0175-z.

13. Derelle, Romain, Hervé Philippe, and John K. Colbourne. 2020. “Broccoli: Combining Phylogenetic and Network Analyses for Orthology Assignment.” Molecular Biology and Evolution 37 (11): 3389–96. 10.1093/molbev/msaa159.

14. Elliott, Glen R. D., and Sally P. Leys. 2010. “Evidence for Glutamate, GABA and NO in Coordinating Behaviour in the Sponge, Ephydatia Muelleri (Demospongiae, Spongillidae).” Journal of Experimental Biology 213 (13): 2310–21. 10.1242/jeb.039859.

15. Ellwanger, Kornelia, Andre Eich, and Michael Nickel. 2007. “GABA and Glutamate Specifically Induce Contractions in the Sponge Tethya Wilhelma.” Journal of Comparative Physiology. A, Neuroethology, Sensory, Neural, and Behavioral Physiology 193 (1): 1–11. 10.1007/s00359-006-0165-y.

16. Ellwanger, Kornelia, and Michael Nickel. 2006. “Neuroactive Substances Specifically Modulate Rhythmic Body Contractions in the Nerveless Metazoon Tethya Wilhelma (Demospongiae, Porifera).” Frontiers in Zoology 3 (1): 7. 10.1186/1742-9994-3-7.

17. Francis, Warren R., Michael Eitel, Sergio Vargas, et al. 2017. “The Genome of the Contractile Demosponge Tethya Wilhelma and the Evolution of Metazoan Neural Signalling Pathways.” Preprint, bioRxiv, April 12. 10.1101/120998.

18. Frickey, Tancred, and Andrei Lupas. 2004. “CLANS: Ag Account Settings for Additional Storage Options. Java Application for Visualizing Protein Families Based on Pairwise Similarity.” Bioinformatics 20 (18): 3702–4. 10.1093/bioinformatics/bth444.

19. Gasnier, Bruno. 2000. “The Loading of Neurotransmitters into Synaptic Vesicles.” Biochimie 82 (4): 327–37. 10.1016/S0300-9084(00)00221-2.

20. Goulty, Matthew, Gaelle Botton-Amiot, Ezio Rosato, Simon G. Sprecher, and Roberto Feuda. 2023. “The Monoaminergic System Is a Bilaterian Innovation.” Nature Communications 14 (1): 3284. 10.1038/s41467-023-39030-2.

21. Hinchliff, Cody E., Stephen A. Smith, James F. Allman, et al. 2015. “Synthesis of Phylogeny and Taxonomy into a Comprehensive Tree of Life.” Proceedings of the National Academy of Sciences 112 (41): 12764–69. 10.1073/pnas.1423041112.

22. Hoang, Diep Thi, Olga Chernomor, Arndt von Haeseler, Bui Quang Minh, and Le Sy Vinh. 2018. “UFBoot2: Improving the Ultrafast Bootstrap Approximation.” Molecular Biology and Evolution 35 (2): 518–22. 10.1093/molbev/msx281.

23. Huang, Ying, Beifang Niu, Ying Gao, Limin Fu, and Weizhong Li. 2010. “CD-HIT Suite: A Web Server for Clustering and Comparing Biological Sequences.” Bioinformatics 26 (5): 680–82. 10.1093/bioinformatics/btq003.

24. Jékely, Gáspár. 2021. “The Chemical Brain Hypothesis for the Origin of Nervous Systems.” Philosophical Transactions of the Royal Society B: Biological Sciences 376 (1821): 20190761. 10.1098/rstb.2019.0761.

25. Jovanovic, Nikola, and Alexander S. Mikheyev. 2019. “Interactive Web-Based Visualization and Sharing of Phylogenetic Trees Using Phylogeny.IO.” Nucleic Acids Research 47 (W1): W266–69. 10.1093/nar/gkz356.

26. Käll, Lukas, Anders Krogh, and Erik L. L. Sonnhammer. 2007. “Advantages of Combined Transmembrane Topology and Signal Peptide Prediction—the Phobius Web Server.” Nucleic Acids Research 35 (suppl_2): W429–32. 10.1093/nar/gkm256.

27. Kanehisa, Minoru, Miho Furumichi, Yoko Sato, Yuriko Matsuura, and Mari Ishiguro-Watanabe. 2025. “KEGG: Biological Systems Database as a Model of the Real World.” Nucleic Acids Research 53 (D1): D672–77. 10.1093/nar/gkae909.

28. Katoh, Kazutaka, John Rozewicki, and Kazunori D. Yamada. 2019. “MAFFT Online Service: Multiple Sequence Alignment, Interactive Sequence Choice and Visualization.” Briefings in Bioinformatics 20 (4): 1160–66. 10.1093/bib/bbx108.

29. Kaupmann, Klemens, Barbara Malitschek, Valerie Schuler, et al. 1998. “GABAB- Receptor Subtypes Assemble into Functional Heteromeric Complexes.” Nature 396 (6712): 683–87. 10.1038/25360.

30. Kristensen, Anders S., Jacob Andersen, Trine N. Jørgensen, et al. 2011. “SLC6 Neurotransmitter Transporters: Structure, Function, and Regulation.” Pharmacological Reviews 63 (3): 585–640. 10.1124/pr.108.000869.

31. Li, Li, Christian J. Stoeckert, and David S. Roos. 2003. “OrthoMCL: Identification of Ortholog Groups for Eukaryotic Genomes.” Genome Research 13 (9): 2178–89. 10.1101/gr.1224503.

32. Li, S. p., H. y. Lee, M. s. Park, J. y. Bahk, B. c. Chung, and M. o. Kim. 2006. “Prenatal GABAB1 and GABAB2 Receptors: Cellular and Subcellular Organelle Localization in Early Fetal Rat Cortical Neurons.” Synapse 60 (8): 557–66. 10.1002/syn.20332.

33. Mao, Chunyou, Cangsong Shen, Chuntao Li, et al. 2020. “Cryo-EM Structures of Inactive and Active GABAB Receptor.” Cell Research 30 (7): 564–73. 10.1038/s41422-020-0350-5.

34. Martin, David L., and Karin Rimvall. 1993. “Regulation of γ-Aminobutyric Acid Synthesis in the Brain.” Journal of Neurochemistry 60 (2): 395–407. 10.1111/j.1471-4159.1993.tb03165.x.

35. Miller, Paul S., and A. Radu Aricescu. 2014. “Crystal Structure of a Human GABAA Receptor.” Nature 512 (7514): 270–75. 10.1038/nature13293.

36. Minh, Bui Quang, Heiko A. Schmidt, Olga Chernomor, et al. 2020. “IQ-TREE 2: New Models and Efficient Methods for Phylogenetic Inference in the Genomic Era.” Molecular Biology and Evolution 37 (5): 1530–34. 10.1093/molbev/msaa015.

37. Morel, Benoit, Alexey M. Kozlov, Alexandros Stamatakis, and Gergely J. Szöllősi. 2020. “GeneRax: A Tool for Species-Tree-Aware Maximum Likelihood-Based Gene Family Tree Inference under Gene Duplication, Transfer, and Loss.” Molecular Biology and Evolution 37 (9): 2763–74. 10.1093/molbev/msaa141.

38. Moroz, Leonid L. 2015. “Convergent Evolution of Neural Systems in Ctenophores.” Journal of Experimental Biology 218 (4): 598–611. 10.1242/jeb.110692.

39. Moroz, Leonid L., Kevin M. Kocot, Mathew R. Citarella, et al. 2014. “The Ctenophore Genome and the Evolutionary Origins of Neural Systems.” Nature 510 (7503): 109–14. 10.1038/nature13400.

40. Moroz, Leonid L., and Andrea B. Kohn. 2016. “Independent Origins of Neurons and Synapses: Insights from Ctenophores.” Philosophical Transactions of the Royal Society B: Biological Sciences 371 (1685): 20150041. 10.1098/rstb.2015.0041.

41. Motiwala, Zenia, Nanda Gowtham Aduri, Hamidreza Shaye, et al. 2022. “Structural Basis of GABA Reuptake Inhibition.” Nature 606 (7915): 820–26. 10.1038/s41586-022-04814-x.

42. Najle, Sebastián R., Xavier Grau-Bové, Anamaria Elek, et al. 2023. “Stepwise Emergence of the Neuronal Gene Expression Program in Early Animal Evolution.” Cell 186 (21): 4676–4693.e29. 10.1016/j.cell.2023.08.027.

43. Patel, Anant B., Robin A. de Graaf, Graeme F. Mason, Douglas L. Rothman, Robert G. Shulman, and Kevin L. Behar. 2005. “The Contribution of GABA to Glutamate/Glutamine Cycling and Energy Metabolism in the Rat Cortex in Vivo.” Proceedings of the National Academy of Sciences 102 (15): 5588–93. 10.1073/pnas.0501703102.

44. Pearson, William R. 2013. “An Introduction to Sequence Similarity (‘homology’) Searching.” Current Protocols in Bioinformatics Chapter 3 (June): 3.1.1-3.1.8. 10.1002/0471250953.bi0301s42.

45. Penel, Simon, Hugo Menet, Théo Tricou, Vincent Daubin, and Eric Tannier. 2022. “Thirdkind: Displaying Phylogenetic Encounters beyond 2-Level Reconciliation.” Bioinformatics 38 (8): 2350–52. 10.1093/bioinformatics/btac062.

46. Pierobon, Paola. 2021. “An Interesting Molecule: γ-Aminobutyric Acid. What Can We Learn from Hydra Polyps?” Brain Sciences 11 (4): 437. 10.3390/brainsci11040437.

47. Pierobon, Paola, Alessandra Concas, Giovanna Santoro, et al. 1995. “Biochemical and Functional Identification of GABA Receptors in *Hydra Vulgaris*.” Life Sciences 56 (18): 1485–97. 10.1016/0024-3205(95)00111-I.

48. Pin, Jean-Philippe, Thierry Galvez, and Laurent Prézeau. 2003. “Evolution, Structure, and Activation Mechanism of Family 3/C G-Protein-Coupled Receptors.” Pharmacology & Therapeutics 98 (3): 325–54. 10.1016/S0163-7258(03)00038-X.

49. Ramoino, Paola, Lorenzo Gallus, Silvio Paluzzi, et al. 2007. “The GABAergic-like System in the Marine Demosponge Chondrilla Nucula.” Microscopy Research and Technique 70 (11): 944–51. 10.1002/jemt.20499.

50. Rost, B. 1999. “Twilight Zone of Protein Sequence Alignments.” Protein Engineering 12 (2): 85–94. 10.1093/protein/12.2.85.

51. Schousboe, Arne, Lasse Kristoffer Bak, and Helle Sønderby Waagepetersen. 2013. “Astrocytic Control of Biosynthesis and Turnover of the Neurotransmitters Glutamate and GABA.” Frontiers in Endocrinology 4 (August). 10.3389/fendo.2013.00102.

52. “Scientific-Inkscape/CITATION.Cff at Main · Burghoff/Scientific-Inkscape.” n.d. Accessed March 5, 2026. https://github.com/burghoff/Scientific-Inkscape/blob/main/CITATION.cff.

53. Seppey, Mathieu, Mosè Manni, and Evgeny M. Zdobnov. 2019. “BUSCO: Assessing Genome Assembly and Annotation Completeness.” In Gene Prediction, edited by Martin Kollmar, vol. 1962. Methods in Molecular Biology. Springer New York. 10.1007/978-1-4939-9173-0_14.

54. Soghomonian, Jean-Jacques, and David L. Martin. 1998. “Two Isoforms of Glutamate Decarboxylase: Why?” Trends in Pharmacological Sciences 19 (12): 500–505. 10.1016/S0165-6147(98)01270-X.

55. Stewart, Gregory D., Laëtitia Comps-Agrar, Lenea Blanc Nørskov-Lauritsen, Jean- Philippe Pin, and Julie Kniazeff. 2018. “Allosteric Interactions between GABAB1 Subunits Control Orthosteric Binding Sites Occupancy within GABAB Oligomers.” Neuropharmacology, A Special Issue Dedicated to Norman G. Bowery, vol. 136 (July): 92–101. 10.1016/j.neuropharm.2017.12.042.

56. “The Ctenophore Genome and the Evolutionary Origins of Neural Systems | Nature.” n.d. Accessed March 6, 2026. https://www.nature.com/articles/nature13400.

57. Tretter, Verena, and Stephen J. Moss. 2008. “GABAA Receptor Dynamics and Constructing GABAergic Synapses.” Frontiers in Molecular Neuroscience 1 (May): 7. 10.3389/neuro.02.007.2008.

58. Tsang, Shui-Ying, Siu-Kin Ng, Zhiwen Xu, and Hong Xue. 2007. “The Evolution of GABAA Receptor–Like Genes.” Molecular Biology and Evolution 24 (2): 599–610. 10.1093/molbev/msl188.

59. Wilkinson, Mark. 2006. “Identifying Stable Reference Taxa for Phylogenetic Nomenclature.” Zoologica Scripta 35 (1): 109–12. 10.1111/j.1463-6409.2005.00213.x.

60. Zaharias, Paul, Frédéric Lemoine, and Olivier Gascuel. 2023. “Robustness of Felsenstein’s Versus Transfer Bootstrap Supports With Respect to Taxon Sampling.” Systematic Biology 72 (6): 1280–95. 10.1093/sysbio/syad052.

61. Zhao, Qinpei, Mantao Xu, and Pasi Fränti. 2008. “Knee Point Detection on Bayesian Information Criterion.” 2008 20th IEEE International Conference on Tools with Artificial Intelligence 2 (November): 431–38. 10.1109/ICTAI.2008.154.

62. Zhou, Y., S. Holmseth, R. Hua, et al. 2012. “The Betaine-GABA Transporter (BGT1, Slc6a12) Is Predominantly Expressed in the Liver and at Lower Levels in the Kidneys and at the Brain Surface.” American Journal of Physiology-Renal Physiology 302 (3): F316–28. 10.1152/ajprenal.00464.2011.

